# An optimized low-dose STZ model rapidly induces diabetic neuropathy in rats

**DOI:** 10.1101/2025.10.02.679000

**Authors:** Glenda Romero-Hernández

## Abstract

**Background:** Diabetic neuropathy is one of the most common complications of diabetes mellitus and remains difficult to study due to the prolonged experimental periods and high mortality frequently associated with conventional streptozotocin (STZ)-induced models. We aimed to develop and characterize a rapid, reproducible, and high-survival rat model of early diabetic neuropathy and to identify molecular alterations associated with disease development.

**Methods:** Male Wistar rats received three intraperitoneal injections of low-dose STZ (30 mg/kg) on alternating days or vehicle control. Blood glucose and body weight were monitored, and a neurophysiological assessment was performed using M-wave and F-wave recordings. To complement functional characterization, dorsal root ganglion (DRG) microarray datasets from STZ-induced diabetic rats and human DRG gene expression samples were analyzed using differential expression, protein-protein interaction, functional enrichment, and receiver operating characteristic (ROC) analyses.

**Results:** STZ-treated animals developed sustained hyperglycemia and significant body weight loss compared with controls. Neurophysiological assessment revealed a marked reduction in F-wave occurrence and prolonged F-wave latency, indicating early peripheral nerve dysfunction. High survival throughout the study. Transcriptomic analysis identified 2,693 differentially expressed genes, with the top 500 enriched in inflammatory, calcium signaling, neuropeptide signaling, and myeloid immune pathways. Network analysis highlighted *TNF, HTR2A, CXCL10*, and *CXCR2* as hub genes. Validation in an independent human DRG dataset demonstrated significant upregulation of *TNF* and *HTR2A*, with strong diagnostic ability (AUC = 0.91 and 0.80, respectively).

**Conclusions:** An optimized low-dose STZ regimen rapidly induces neurophysiological features of diabetic neuropathy within four weeks while maintaining high animal survival. This model provides a practical platform for investigating early pathogenic mechanisms and evaluating therapeutic interventions. Integrative transcriptomic analyses further identify *TNF* and *HTR2A* as candidate biomarkers with translational relevance for diabetic neuropathy.

## Main

Diabetes mellitus is a rising global health challenge, with an estimated 3.4 million deaths in 2024 and projections suggesting that the number of affected individuals will exceed 800 million by 2050^1^. Diabetic neuropathy is among the most common and debilitating complications of diabetes, contributing to pain, falls, ulceration, infection, autonomic dysfunction and limb amputation^2^. Chronic hyperglycaemia promotes multiple pathogenic mechanisms, including oxidative stress, advanced glycation end-product formation and inflammatory signalling, which impair axons, Schwann cells and the microvasculature, ultimately compromising peripheral nerve function^3^.

Streptozotocin (STZ)-induced diabetes remains one of the most widely used experimental models for studying diabetic neuropathy. However, conventional STZ regimens are frequently associated with high mortality, severe metabolic deterioration and inter-study variability, limiting their utility for longitudinal investigations and reducing experimental reproducibility^4^. Furthermore, many studies focus on late-stage manifestations of neuropathy, often assessed after eight or more weeks of diabetes induction, whereas early neuropathic alterations remain poorly characterized. Therefore, there is a need for new experimental models that combine reliable diabetes induction, reproducible neuropathic features and high animal survival within a relatively short experimental timeframe.

A critical aspect in the characterization of diabetic neuropathy models is the identification of objective and clinically relevant biomarkers capable of detecting early nerve dysfunction. Electrophysiological assessments provide quantitative measures of peripheral nerve impairment and can reveal neuropathic alterations before the onset of severe clinical manifestations^5^. Among these, F-waves evaluate proximal motor pathway integrity and have been shown to be sensitive, reproducible and clinically translatable indicators of peripheral nerve dysfunction^5^. Consequently, F-wave analysis represents a valuable approach for assessing early neuropathic changes in both experimental models and clinical settings. Building upon these findings, the present study focused on the early phase of disease development and aimed to establish and characterize an optimized low-dose STZ-induced rat model of diabetic neuropathy. Specifically, we evaluated whether this protocol could maintain hallmark metabolic alterations while enabling the detection of early neurophysiological abnormalities within four weeks of diabetes induction without compromising animal survival. To complement the functional assessment provided by F-wave recordings, we also analyzed dorsal root ganglion (DRG) gene expression datasets from rats and humans to investigate molecular alterations associated with the sensory component of diabetic neuropathy. Together, these approaches provide an integrated characterization of the model at both the neurophysiological and molecular levels.

## Results

### Experimental design of the STZ-induced diabetic neuropathy model

To establish a short-term model of diabetic neuropathy, diabetes was induced in adult rats by administering three intraperitoneal injections of streptozotocin (STZ: 30 mg/kg) on alternating days (days 1, 3, and 5). Body weight and blood glucose levels were monitored at baseline and at the study endpoint to assess the metabolic effects of STZ treatment. Four weeks after diabetes induction, neurophysiological evaluation was performed using F-wave recordings to assess peripheral nerve abnormalities associated with diabetic neuropathy, and blood glucose levels were measured from tail vein samples. The overall experimental workflow is summarized in **Fig. 1**.

**Fig. 1:**
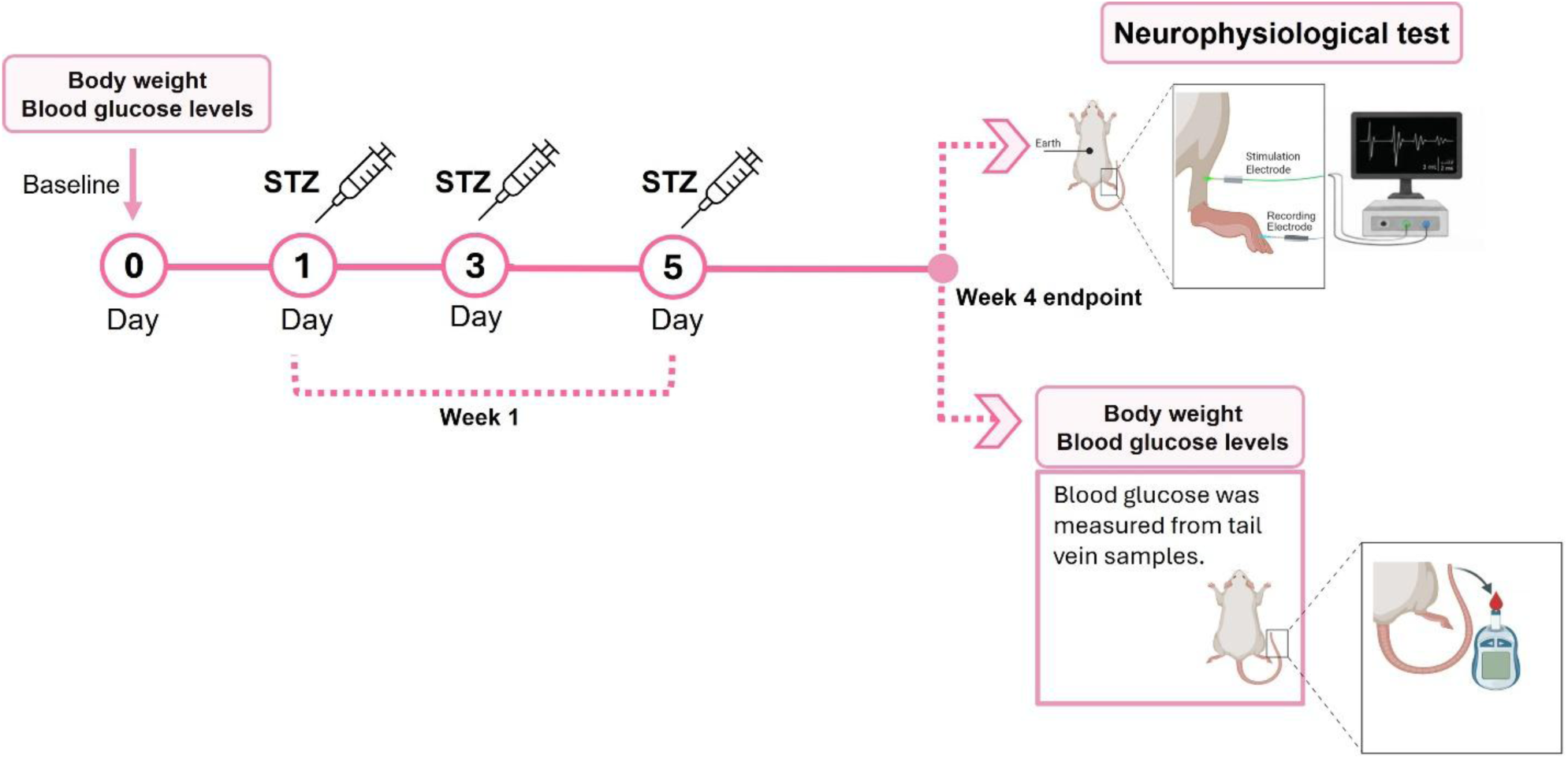
Experimental design of the STZ-induced diabetic neuropathy model. Overview of the experimental workflow. Body weight and blood glucose levels were monitored at baseline and at week 4. Diabetes induction was generated by three STZ (30 mg/kg) injections administered on alternating days (days 1, 3, and 5). At the week 4, neurophysiological assessment was conducted using F-wave recordings, and blood glucose levels were measured from tail vein samples.

### Validation of diabetes induction in a short-term neuropathy model

At baseline, blood glucose levels were comparable between control and STZ-treated rats, with all animals displaying values within the normal physiological range of 4-6 mmol/L (**Fig. 2a**). Four weeks after STZ administration, diabetic animals displayed marked hyperglycemia, whereas glucose levels remained unchanged in controls. Blood glucose concentrations were significantly higher in the STZ-treated group than in control animals at the study endpoint (**Fig. 2a**). Together, these findings demonstrated the establishment of a diabetic phenotype within the four-week study period.

**Fig. 2:**
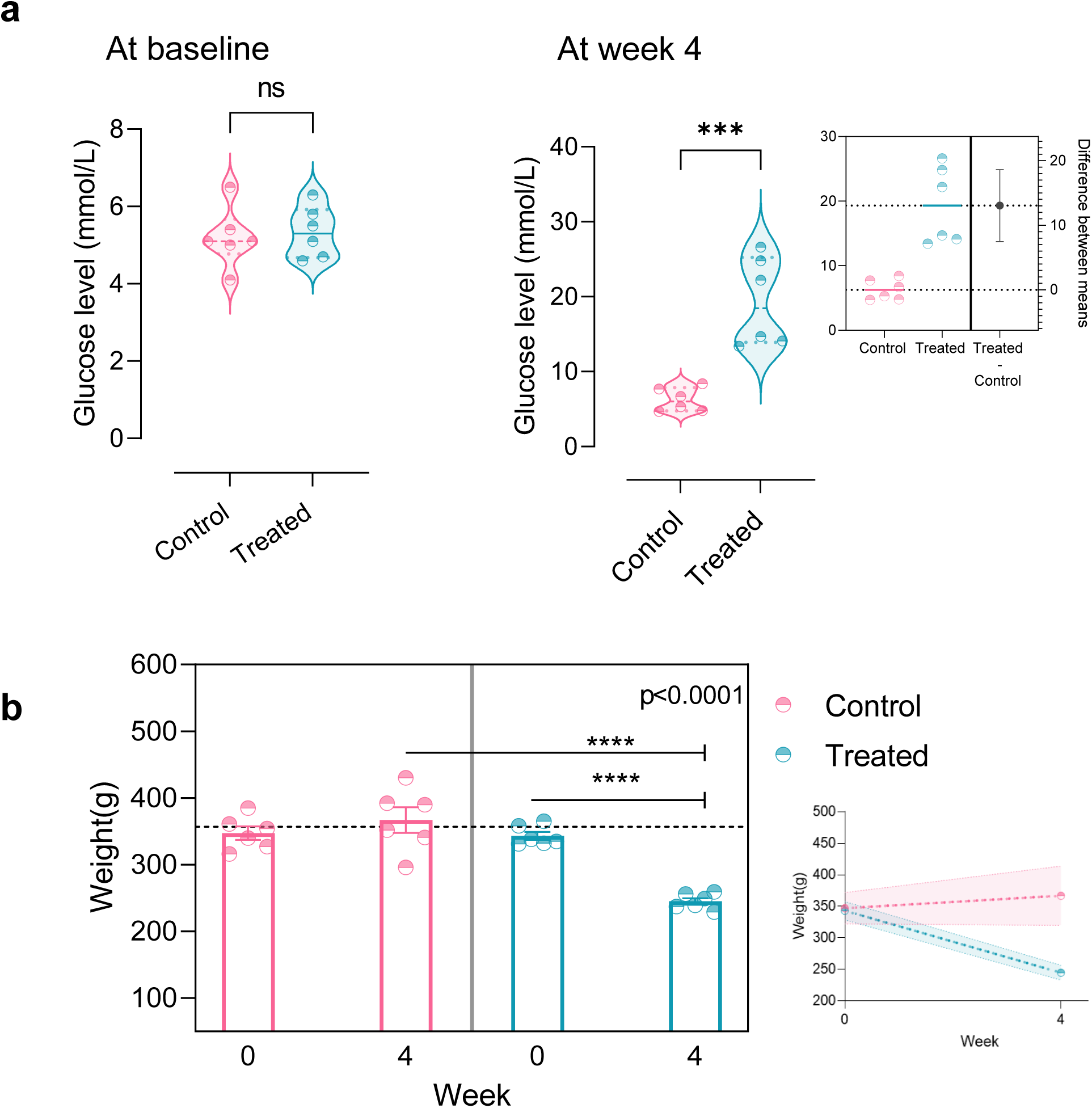
Validation of diabetes induction in the STZ-treated rat model. a, Blood glucose levels measured in control and STZ-treated animals at baseline and at week 4. The right panel shows the difference between group means at week 4 together with the corresponding 95% confidence interval. b, Body weight measurements obtained at baseline and week 4 in control and STZ-treated animals. Bars represent mean values and points represent individual animals. Statistical comparisons were performed using unpaired or paired Student’s t-tests, as indicated in the figures. Data are presented as mean ± SEM. ns, not significant; ***P < 0.001; ****P < 0.0001.

In parallel, body weight was similar between groups at baseline (**Fig. 2b**). Over the 4-week experimental period, STZ-treated rats showed a progressive reduction in body weight, whereas control animals maintained stable weights. Within-group analysis revealed a significant decrease in body weight in diabetic animals between baseline and week 4 (**Fig. 2b**). Consequently, STZ-treated rats displayed lower body weights than controls at the study endpoint, consistent with the metabolic consequences of experimental diabetes.

### Early neurophysiological alterations in STZ-treated rats

To determine whether STZ-induced diabetes was associated with peripheral nerve dysfunction, neurophysiological assessment was performed four weeks after diabetes induction using F-wave recordings. Representative M-wave and F-wave recordings obtained from three animals per group revealed clear differences between control and STZ-treated rats (**Fig. 3a**). Quantitative analysis demonstrated a marked reduction in F-wave occurrence in STZ-treated animals compared with controls (**Fig. 3b**), indicating peripheral nerve dysfunction.

**Fig. 3:**
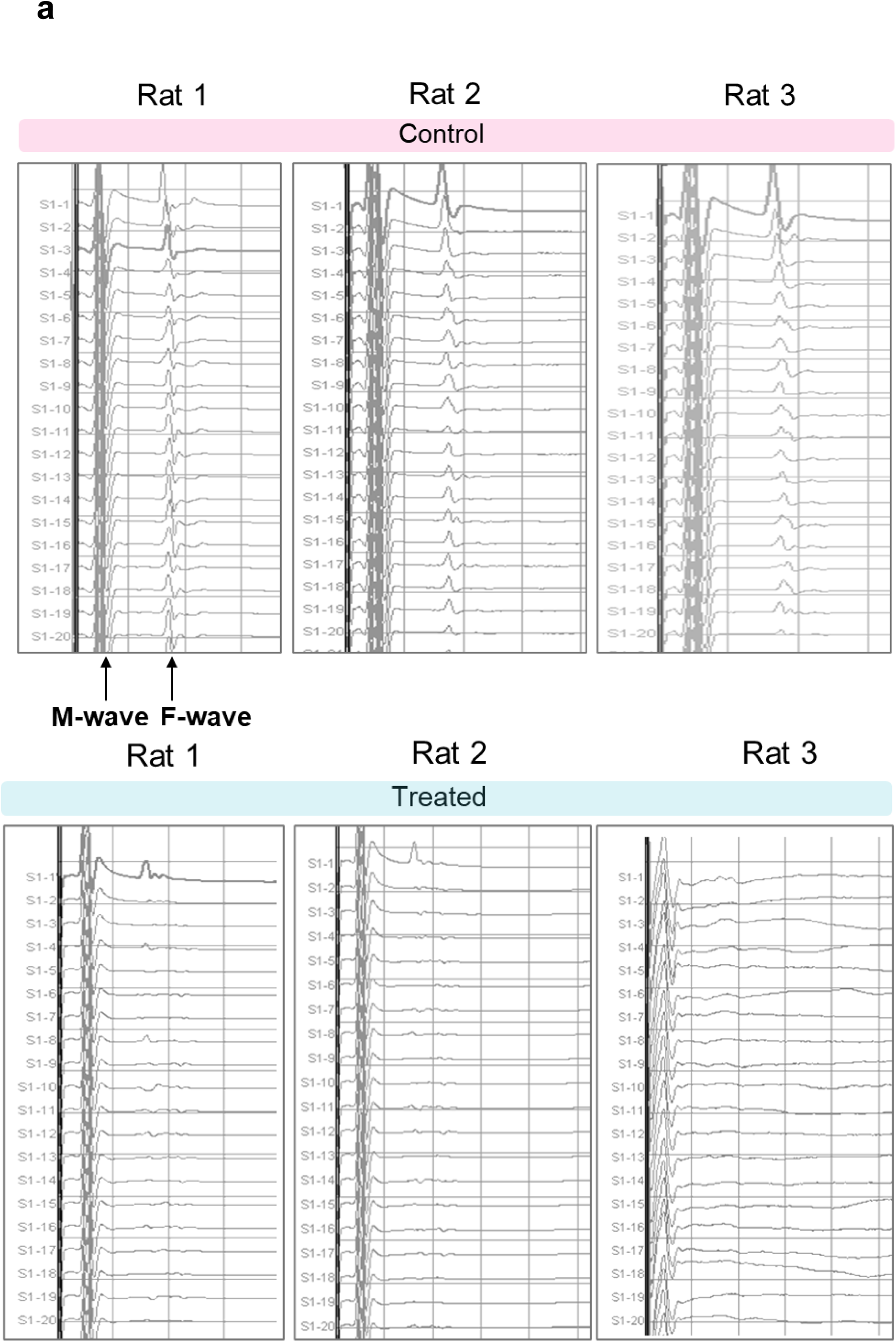
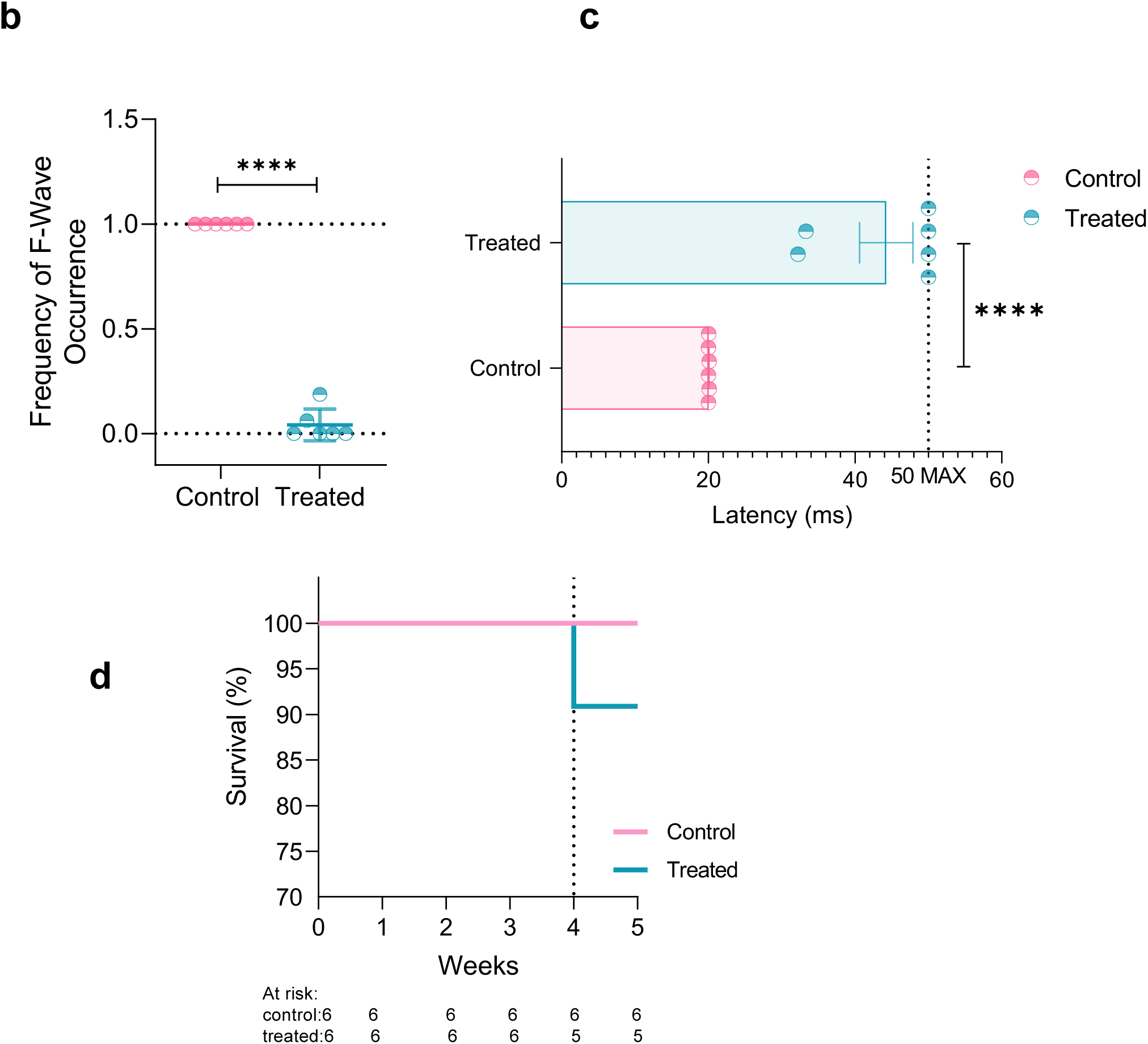
Neurophysiological assessment and survival of control and STZ-treated rats. a, Representative recordings of M-waves and F-waves obtained from three control and three STZ-treated animals. b, Frequency of F-wave occurrence following repetitive stimulation in control and STZ-treated rats. c, F-wave latency measurements in control and STZ-treated animals, with 50 ms being the maximum latency. d, Kaplan-Meier survival curves for control and STZ-treated rats throughout the 4-week experimental period using the log-rank test. Numbers at risk are shown below the x-axis. Statistical comparisons were performed using unpaired Student’s t-tests, and data are presented as mean ± SEM. ****P < 0.0001.

Moreover, F-wave latency was significantly prolonged in STZ-treated animals relative to controls (**Fig. 3c**). Of note, recordings in which no F-wave was detected within the recording window were assigned the maximum measurable latency value (50 ms). Most diabetic animals reached this threshold, indicating marked impairment of F-wave generation and peripheral nerve conduction. Together, these findings demonstrate that neurophysiological alterations were detectable within four weeks of STZ administration and support the development of early diabetic neuropathy in this model.

On the other hand, survival remained high throughout the study period in both groups. All control animals survived until the experimental endpoint, whereas one death occurred in the STZ-treated group during week 4, resulting in a survival rate of 83.3% compared with 100% in controls (**Fig. 3d**). Kaplan-Meier analysis revealed no significant differences in survival between groups during the 4-week experimental period (log-rank test, p = 0.31). Notably, the death occurred during anesthesia following electrophysiological assessment and was therefore not attributed to the diabetic condition.

### Transcriptomic alterations associated with diabetic neuropathy

To complement the neurophysiological characterization of our diabetic neuropathy model, we analyzed a DRG gene expression dataset from STZ-induced diabetic rats collected four weeks after diabetes induction. PCA analysis revealed clear separation between diabetic and control samples, with PC1 accounting for 82% of the total variance. Consistently, hierarchical clustering identified two distinct clusters corresponding to diabetic and control animals, reflecting differences in gene expression profiles between the two groups (**Fig. 4a**). From the differential expression analysis, 2,693 genes were identified as significantly altered in diabetic rats (adjusted p < 0.05, |log2 fold change| > 1.5). Among these, 88.64% were upregulated and 11.36% were downregulated relative to controls (**Fig. 4b**). To identify key molecular regulators associated with diabetic neuropathy, a PPI network was constructed using the top 500 differentially expressed genes. Network analysis revealed a subset of highly interconnected nodes and identified nine hub proteins (dark pink) with the highest degree of connectivity within the network (**Fig. 4c**). Moreover, proteins included in the interaction network were overrepresented in several pathways associated with neuronal dysfunction. Functional enrichment analysis revealed that upregulated genes were enriched in *calcium signaling, adenylate cyclase-activating signaling, phospholipase C-activating signaling, inflammatory response, neuropeptide signaling,* and *myeloid leukocyte-mediated immunity* pathways (**Fig. 4d**), suggesting activation of inflammatory and neuronal signaling processes in diabetic animals. In contrast, downregulated genes were enriched in pathways related to *neuron differentiation, phosphotransferase activity, nucleic acid metabolic processes,* and *positive regulation of transcription by RNA polymerase II* (**Fig. 4d**), suggesting suppression of gene regulatory and neuronal maintenance programs.

**Fig. 4:**
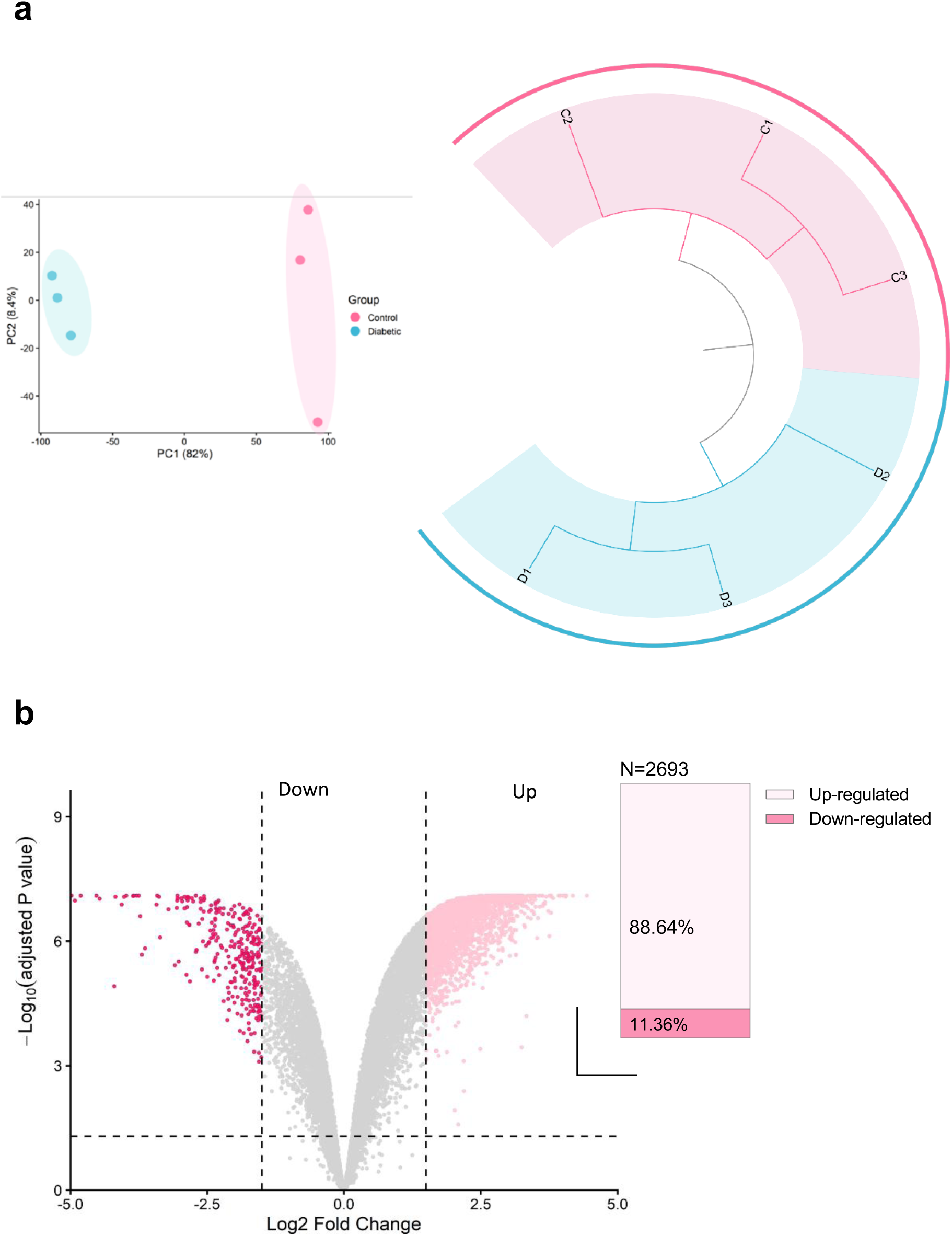
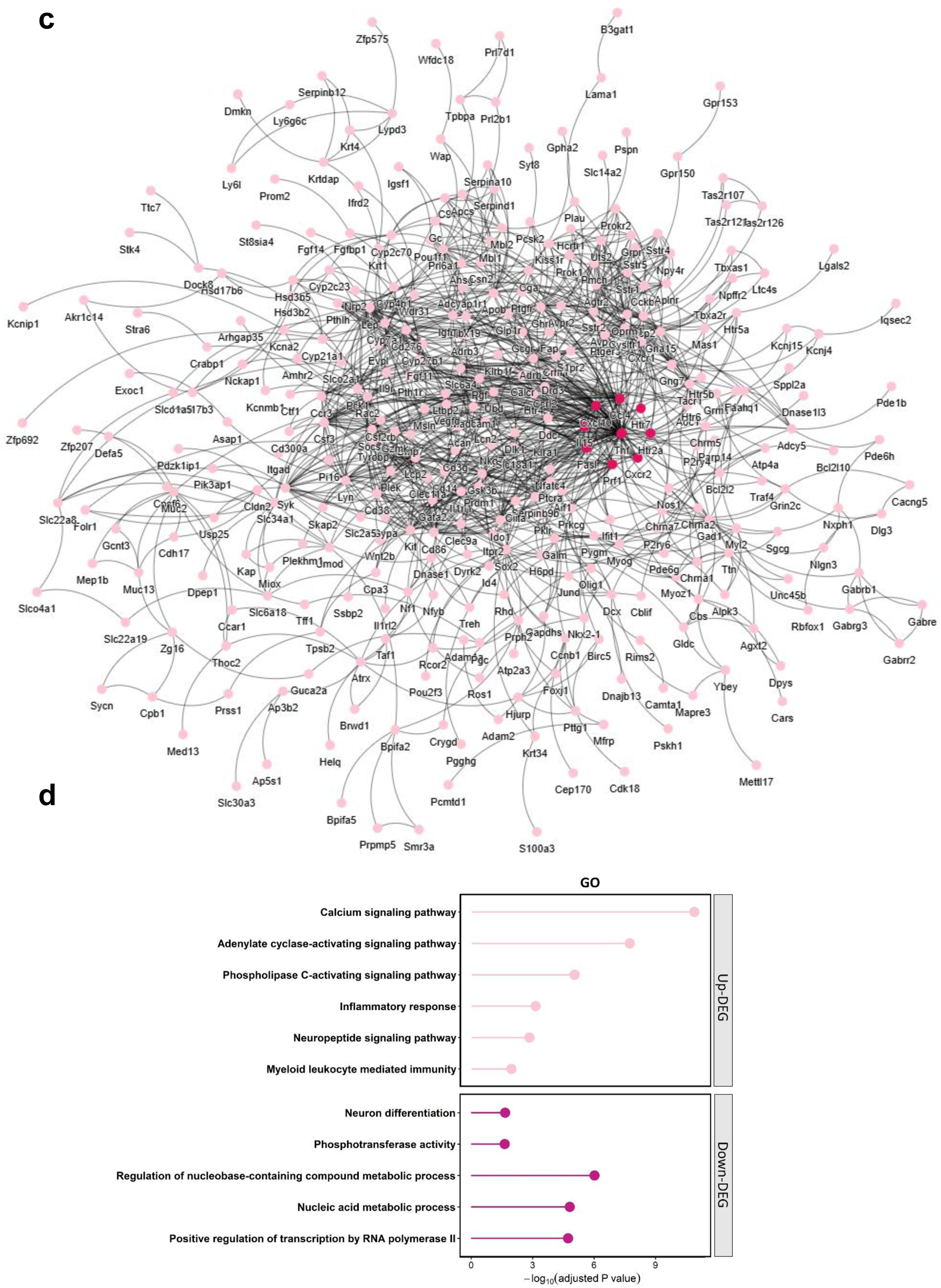
Identification and functional analysis of differentially expressed genes in diabetic rats. a, Principal component analysis (PCA) and hierarchical clustering of gene expression profiles from control and diabetic dorsal root ganglion samples. b, Volcano plot showing differential gene expression between diabetic and control groups. Dashed lines indicate the significance thresholds (|log2 fold change| > 1 and adjusted p value < 0.05). The right panel summarizes the total number (N) and proportion of upregulated and downregulated genes identified in the analysis. c, Protein-protein interaction network generated from the top 500 differentially expressed genes. Nodes represent genes and edges represent known interactions. Highlighted nodes (dark pink) indicate highly connected genes within the network. d, Functional enrichment analysis of differentially expressed genes included in the network. Enriched biological pathways are shown separately for upregulated and downregulated gene sets.

To prioritize candidate biomarkers associated with early diabetic neuropathy, among the nine hub genes identified by network analysis, *TNF*, *HTR2A*, *CXCL10*, and *CXCR2* were selected for further validation. These genes combined high connectivity within the PPI network with recurrent representation across multiple significantly enriched pathways among upregulated genes, suggesting a central role in the molecular alterations associated with diabetic neuropathy. Notably, all four genes showed log2 fold-change values greater than 2, corresponding to more than 4-fold increases in expression relative to controls, highlighting their strong transcriptional activation in diabetic animals and their potential for reproducible detection across independent datasets (**Fig. 5a**).

**Fig. 5:**
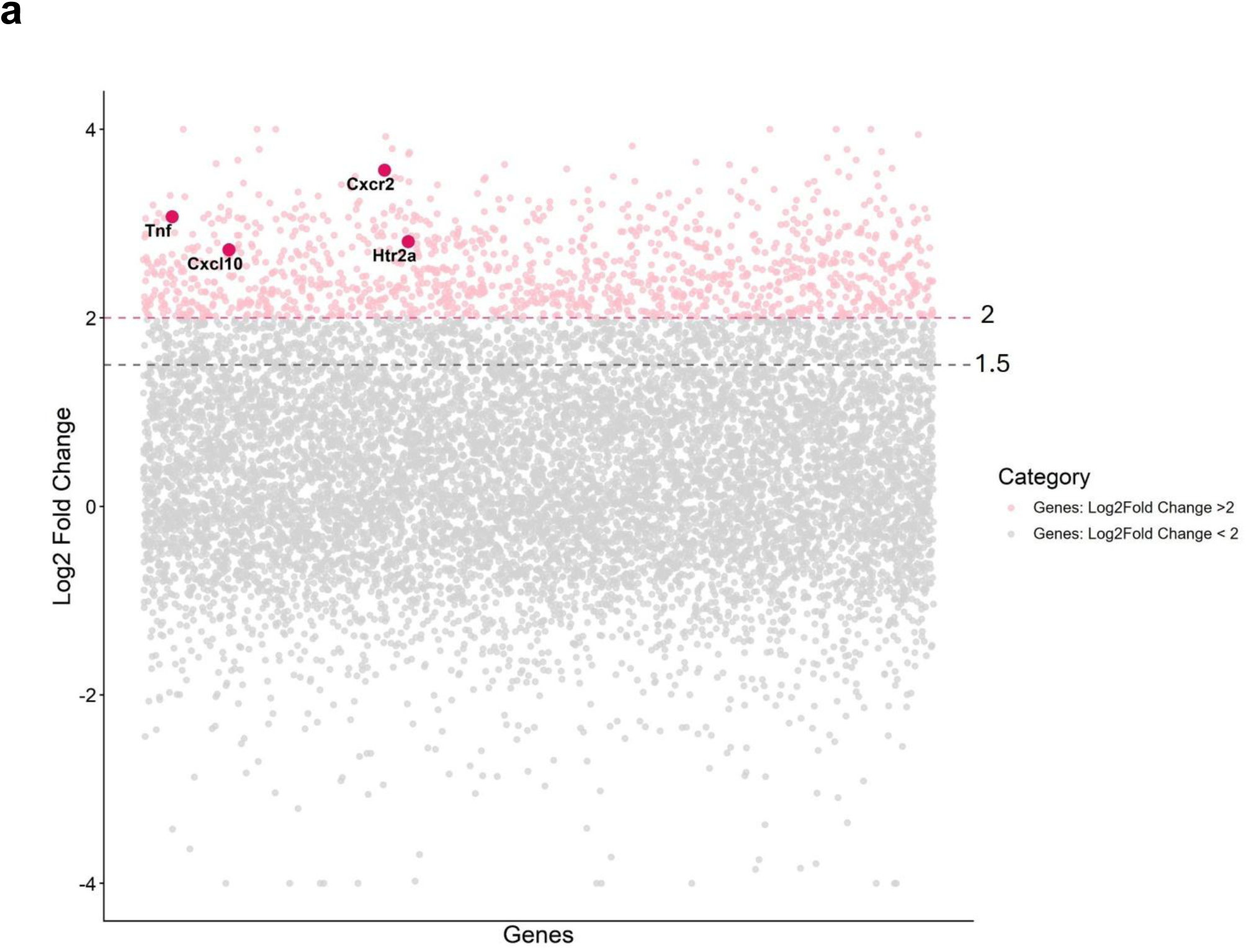
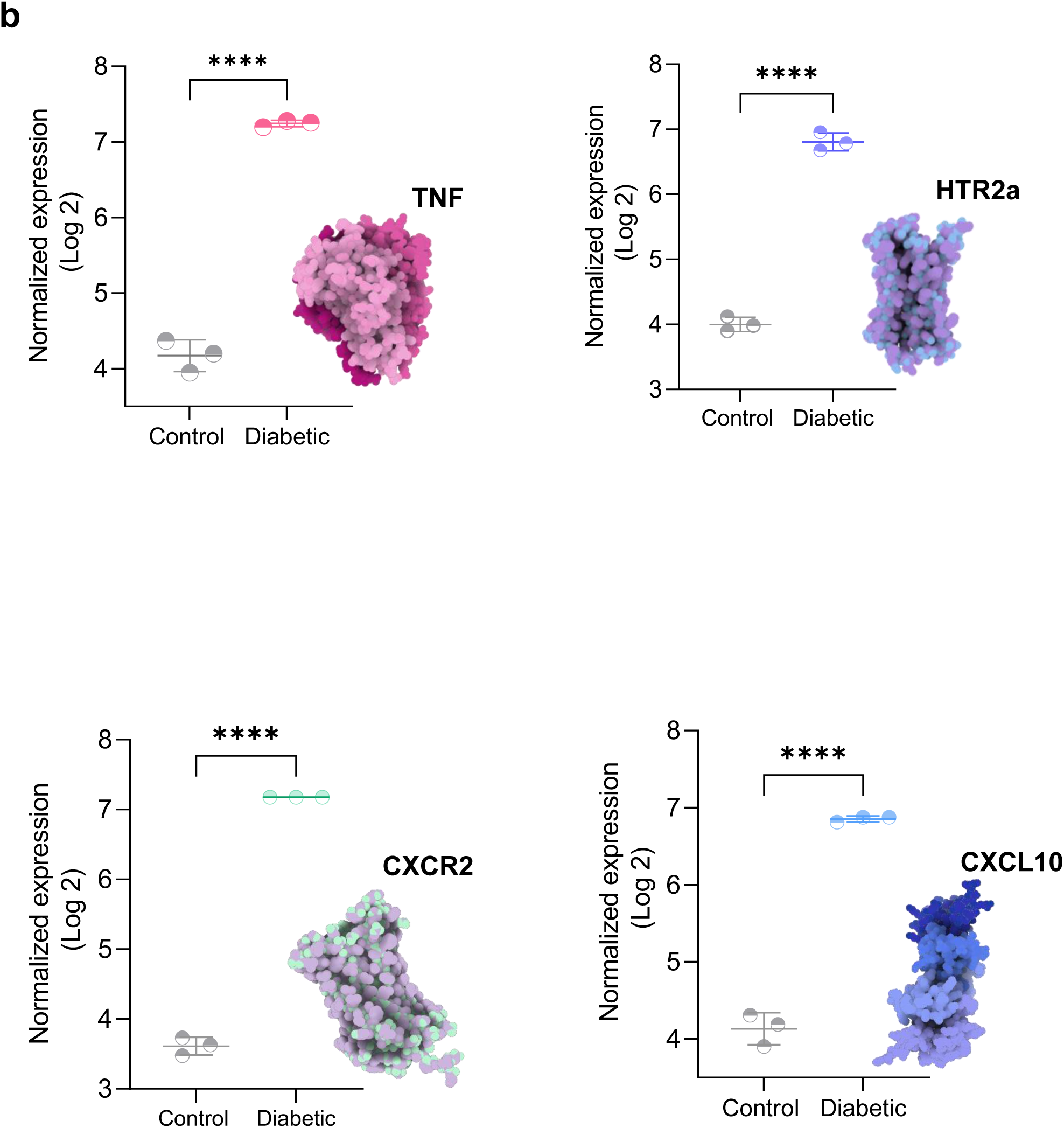
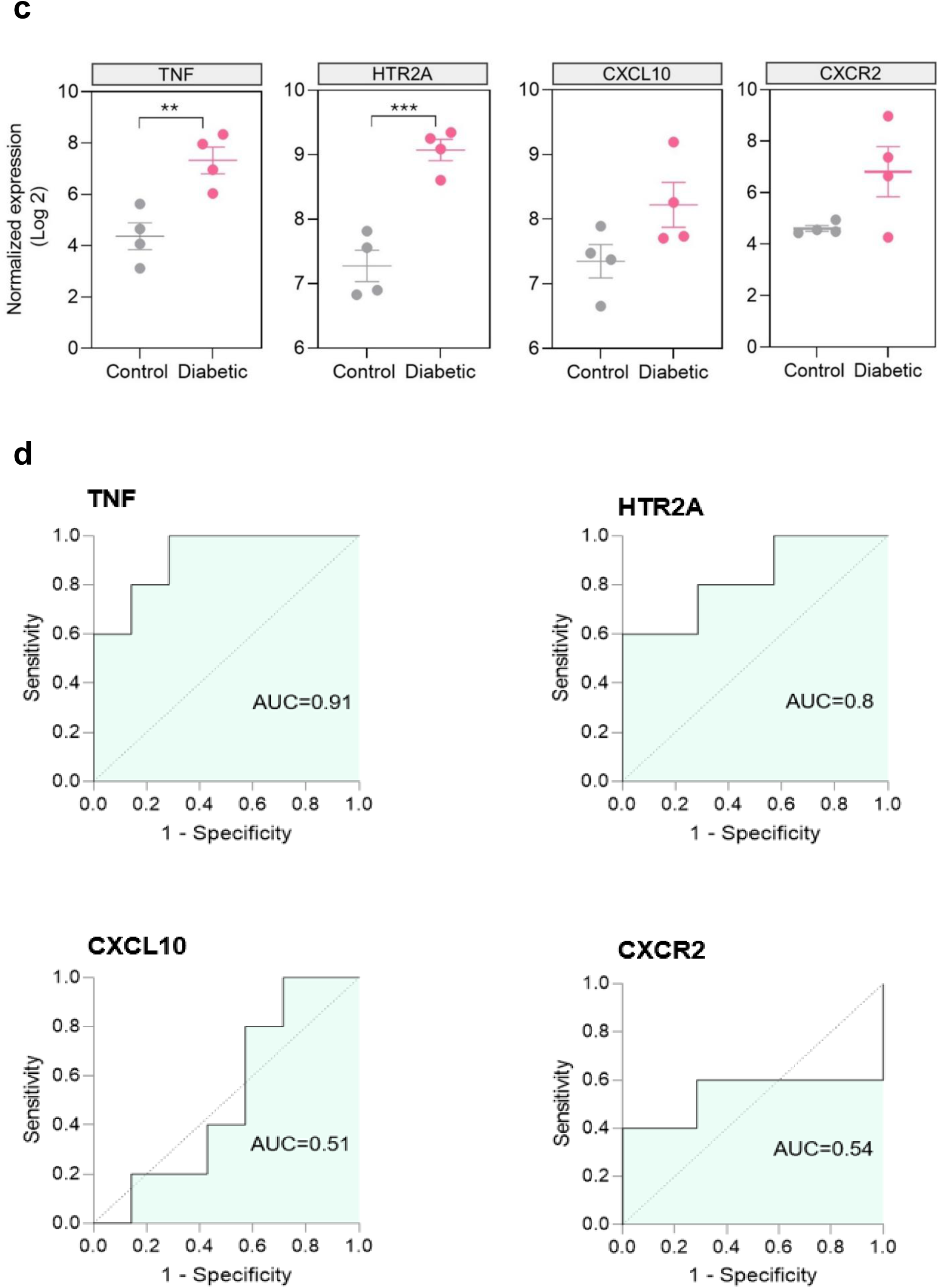
Expression analysis of selected hub genes. a, Differential gene expression analysis highlighting *TNF, HTR2A, CXCL10 and CXCR2* among all genes in diabetic rats. b, Normalized expression levels (log2) of TNF, HTR2A, CXCL10 and CXCR2 in control and diabetic rats. Representative protein structures corresponding to each gene product are displayed alongside the expression plots. c, Expression levels of *TNF, HTR2A, CXCL10* and *CXCR2* shown individually from a human dataset. d, ROC curve analysis evaluating the discriminatory power of the selected hub genes in an independent human cohort. Statistical comparisons were performed using unpaired Student’s t-tests, and data are presented as mean ± SEM. *p<0.05, **p < 0.01, ***p < 0.001.

Expression analysis confirmed significant differences in the expression of *TNF, HTR2A, CXCL10*, and *CXCR2* in diabetic animals compared with controls (**Fig. 5b**). The clear separation observed between diabetic and control samples, together with the low within-group variability, supported the reproducibility of these transcriptional alterations.

### Diagnostic evaluation of candidate hub genes in an independent human dataset

To assess their translational relevance as potential biomarkers of early neuronal dysfunction in diabetic patients, the expression patterns of these genes were evaluated in an independent human DRG dataset. *TNF* and *HTR2A* remained significantly elevated in diabetic samples, whereas *CXCL10* and *CXCR2* showed no statistically significant differences between groups (**Fig. 5c**). ROC curve analysis was subsequently performed to evaluate the discriminatory power of these four genes and further confirm their potential as biomarkers of early neuronal dysfunction. *TNF* demonstrated the highest diagnostic performance (AUC = 0.91), followed by *HTR2A* (AUC = 0.80). In contrast, *CXCL10* and *CXCR2* showed no discriminative ability, with AUC values of 0.51 and 0.54, respectively (**Fig. 5d**). Thus, our findings identify *TNF* and *HTR2A* as the most promising candidate biomarkers for early neuropathy in diabetic patients.

## Discussion

Our study describes a short-term STZ-induced model of diabetic neuropathy characterized by sustained hyperglycemia, weight loss, and neurophysiological abnormalities within four weeks of diabetes induction. These findings demonstrate that the proposed protocol successfully reproduces both the metabolic and neurological features of diabetic neuropathy within a relatively short experimental timeframe. In addition, gene expression analysis revealed molecular alterations associated with neuroinflammatory and neuronal signaling pathways that may underlie the early development of diabetic neuropathy.

Experimental models of diabetes remain essential tools for investigating disease mechanisms and evaluating novel therapeutic strategies while reducing the need for ethically challenging studies in human subjects ^6^. An ideal animal model should reproduce not only the metabolic abnormalities associated with diabetes but also the progression of clinically relevant complications ^7^. Among the available experimental approaches, STZ-induced diabetes remains one of the most widely used and reproducible models because it selectively targets pancreatic β-cells and induces sustained hyperglycemia ^8^. Following uptake through the GLUT2 transporter, STZ causes deoxyribonucleic acid alkylation, oxidative stress, and disruption of cellular metabolism, ultimately leading to insulin deficiency and diabetic pathology ^9^. As a result, STZ-based models have been extensively used to investigate the development of diabetes. However, the onset and severity of neuropathic manifestations vary substantially depending on STZ dose, administration schedule, and duration of follow-up. In many studies, neuropathic phenotypes are characterized after 6-12 weeks or longer, whereas our model detected significant neurophysiological abnormalities within only four weeks after diabetes induction ^6,10,11^. The rapid onset of F-wave abnormalities observed in the present study suggests that this protocol captures early functional alterations of peripheral nerves before the development of more advanced neuropathic complications. Importantly, survival remained high throughout the study period, with only a single mortality event occurring during anesthesia following electrophysiological assessment rather than as a direct consequence of diabetes. This represents a potential advantage over more aggressive STZ protocols, which may be associated with greater systemic toxicity, severe metabolic deterioration, and increased mortality.

Neurophysiological assessment revealed a marked reduction in F-wave occurrence, together with prolonged F-wave latency in diabetic animals. F-wave recordings provide information regarding the functional integrity of motor neurons and proximal nerve segments and are widely used to detect abnormalities in peripheral nerve conduction. The observation that most diabetic animals reached the maximum measurable latency threshold indicates substantial impairment of neural excitability and signal propagation despite the relatively short duration of diabetes. These findings suggest that nerve dysfunction develops early during diabetic neuropathy.

F-wave recordings provide information regarding the functional integrity of motor neurons and proximal nerve segments and are widely used to detect abnormalities in peripheral nerve conduction ^12^. Importantly, because F-waves assess conduction along the entire length of the motor pathway, they are considered parameters for detecting early neuropathic changes. Several clinical studies have reported prolonged F-wave latency and reduced F-wave persistence in patients with diabetic neuropathy, even in cases where routine nerve conduction studies remain within normal ranges, supporting their utility as early indicators of neural dysfunction ^5^. Notably, previous studies have identified F-wave latency as one of the most sensitive electrophysiological parameters for detecting diabetic nerve pathology and subclinical neuropathy ^5,13–15^. Thus, the reduction in F-wave occurrence and prolongation of latency observed in our study indicate that this short-term STZ protocol successfully reproduces early functional abnormalities characteristic of diabetic neuropathy. The ability to detect these alterations within four weeks further supports the utility of this model for investigating mechanisms involved in the initial stages of disease progression before the development of more advanced structural nerve damage.

To further investigate molecular mechanisms associated with these neurophysiological changes, we analyzed an independent DRG transcriptomic dataset obtained four weeks after STZ administration in rats. Gene expression profiling revealed clear differences between diabetic and control animals, with a high number of differentially expressed genes. Moreover, most of the top 500 differentially expressed genes were connected within a biological interaction network, suggesting coordinated molecular responses associated with diabetic neuropathy. In this regard, functional enrichment analysis demonstrated activation of inflammatory pathways, neuropeptide signaling, calcium signaling, phospholipase C signaling, and myeloid leukocyte-mediated immunity ^16^. These results are consistent with growing evidence indicating that diabetic neuropathy is not solely a metabolic disorder but also involves a substantial neuroinflammatory component characterized by immune cell activation, cytokine production, and dysregulation of neuronal signaling pathways ^3,17^. In contrast, downregulated genes were enriched in pathways related to neuron differentiation, nucleic acid metabolism, and transcriptional regulation, suggesting impairment of neuronal maintenance and gene regulatory programs during the early stages of diabetic neuropathy. Together, these results support the concept that diabetic neuropathy involves both activation of inflammatory signaling and disruption of pathways required for normal neuronal function.

Network analysis identified several highly connected hub genes, where *TNF, HTR2A, CXCL10,* and *CXCR2* were recurrently represented across enriched pathways. Among these candidates, *TNF* emerged as the strongest biomarker. *TNF* is a central pro-inflammatory cytokine that has been repeatedly implicated in the pathogenesis of diabetic neuropathy ^18^. Elevated TNF levels have been reported in patients with diabetic neuropathy and have been associated with reduced nerve conduction velocity, increased pain perception, and enhanced neuronal excitability ^19^. Experimental studies further demonstrate that *TNF and CXCL10* contribute to neuroinflammation through activation of immune cells and sensitization of peripheral neurons ^19–21^. Consistent with these observations, *TNF* was strongly upregulated in diabetic rats, remained significantly elevated in the independent human dataset, and demonstrated the highest diagnostic performance among the genes evaluated.

*HTR2A* also emerged as a promising candidate biomarker. Although less studied than *TNF* in diabetic neuropathy, serotonergic signaling plays an important role in pain modulation, sensory processing, and neuronal excitability. Previous studies have implicated serotonin receptors, including HTR2A-related pathways, in neuropathic pain mechanisms and altered nociceptive processing in diabetic neuropathy models ^22,23^. The persistence of HTR2A overexpression in the human cohort, together with its favorable discriminatory ability, suggests that serotonergic dysregulation may contribute to early neuronal dysfunction associated with diabetic neuropathy. These findings are particularly interesting because they identify a candidate biomarker that extends beyond classical inflammatory mechanisms and may reflect alterations in neuronal signaling pathways during the early stages of disease development.

Several considerations should be taken into account when interpreting these findings. The transcriptomic analysis was performed using microarray data from DRG tissue, providing a comprehensive overview of molecular alterations associated with diabetic neuropathy. However, future studies using single-cell or spatial transcriptomic approaches could further clarify the specific cellular populations contributing to the observed expression patterns. In addition, although the independent human dataset supported the translational relevance of *TNF* and *HTR2A*, validation in larger patient cohorts would help to further assess their potential clinical utility.

The present study was primarily designed to establish the presence of diabetes and verify the development of neuropathy through neurophysiological assessment. Given that nerve conduction studies represent sensitive and minimally invasive approaches for evaluating peripheral nerve dysfunction, future investigations could further characterize this model through histopathological analyses of affected neural tissues. Such studies would provide complementary structural information and improve understanding of the relationship between early neurophysiological abnormalities and tissue-level alterations. Furthermore, the low mortality observed in this work supports the feasibility of extending the duration of this model, allowing evaluation of disease progression and providing a platform for testing candidate therapeutic interventions after neuropathy has been established. In addition, future experiments incorporating insulin or other glucose-lowering therapies may better reflect the clinical setting, where patients with diabetic neuropathy typically receive standard treatments for glycemic control while undergoing evaluation of potential neuroprotective or anti-neuropathic therapies.

In conclusion, we optimized a rapid and reproducible low-dose STZ model that develops key neurophysiological features of diabetic neuropathy within four weeks while maintaining high animal survival. These findings indicate that the proposed protocol provides an efficient and practical approach for studying early diabetic neuropathy, reducing the prolonged experimental periods frequently required in conventional STZ-induced models while preserving animal welfare. Similarly, integrative transcriptomic analysis revealed molecular alterations associated with inflammatory and neuronal signaling pathways, supporting the concept that neuroinflammation and neuronal dysfunction are early events in the development of diabetic neuropathy. Among the identified hub genes, *TNF* and *HTR2A* emerged as the most promising candidate biomarkers, demonstrating translational relevance in human samples.

Collectively, this model provides a useful platform for investigating the early mechanisms underlying diabetic neuropathy and for evaluating novel therapeutic strategies targeting neuroinflammatory and neuronal signaling pathways.

## Materials and Methods

### Animals

Male Wistar rats (6-8 weeks of age, approximately 300 g body weight) were housed under standard laboratory conditions. Animals were acclimated for one week before the start of the experiments under a 12:12 hours light-dark cycle at 24-26 °C and 60% relative humidity, with ad libitum access to food and water. Rats were randomly assigned to either the control group (n = 6) or the STZ-treated group (n = 6).

### Induction of diabetes mellitus

STZ (Sigma-Aldrich, UK) was stored at -20 °C and freshly dissolved in citrate-phosphate buffer (pH 4.5) immediately before administration. Diabetes was induced by three intraperitoneal injections of STZ at low dose of 30 mg/kg. Control animals received equivalent volumes of citrate-phosphate buffer without STZ. Animal handling and anesthesia procedures were performed as previously described^12^.

### Validation of the diabetic neuropathy model

Blood glucose levels and body weight were used to assess diabetes induction and model establishment. To minimize animal stress, blood glucose measurements were performed under non-fasting conditions. Blood glucose levels and body weight were recorded at baseline and at the 4-week endpoint. For each measurement, the tail tip was cleaned, and a small blood sample was collected for glucose determination using a conventional glucometer (SUMA, Cuba).

Electrophysiological recordings were performed at week 4 using a Neuronic-5 system (Neuronic S.A., Cuba). Animals were anesthetized with chloral hydrate (400 mg/kg, intraperitoneal) and positioned in dorsal recumbency for electrode placement. F-waves were elicited using two surface clip electrodes placed on either side of the distal tibia, proximal to the ankle joint. Responses were recorded using a surface clip electrode positioned on the dorsal aspect of the foot as the active electrode, a needle electrode inserted into the interdigital membrane as the reference electrode, and a needle electrode placed in the ventral skin as the ground. Stimulation was delivered at a frequency of 10 Hz, and a train of 20 supramaximal stimuli was applied to each animal. F-wave frequency was calculated as the proportion of detectable F-wave responses within the stimulus train. For latency analyses, recordings in which no F-wave was detected within the expected response window were assigned the maximum latency value of the recording window (50 ms) for statistical analysis.

### Bioinformatic analysis of dorsal root ganglion gene expression

To complement the neurophysiological characterization of our diabetic neuropathy model, a DRG gene expression dataset was analyzed. rat gene expression data were obtained from the ArrayExpress database (accession E-MEXP-515), generated using Affymetrix GeneChip oligonucleotide microarrays. The dataset included DRG samples from control and STZ-induced diabetic rats collected at 4 weeks after diabetes induction^24^.

Principal component analysis (PCA) was performed in R using the *prcomp* function from the stats package^25^. Unsupervised hierarchical clustering based on Euclidean distance and Ward’s linkage method was performed using the *hclust* function and visualized with the *ggtree* package^26^. These analyses were used to assess global gene expression patterns and sample clustering.

Differential expression analysis between diabetic and control samples was performed using the *limma* package^27^. Statistical significance was assessed using moderated t-tests, and p-values were adjusted for multiple testing using the Benjamini-Hochberg method^28^. Genes with a |log2 fold change| > 1.5 and an adjusted P value < 0.05 were considered differentially expressed (DE).

### Protein interaction network and functional enrichment analysis

Protein-protein interaction (PPI) analysis was performed using the Search Tool for the Retrieval of Interacting Genes/Proteins (STRING) database to investigate interactions among proteins encoded by top 500 DE genes^29^. The resulting network was analyzed to identify hub genes based on network connectivity. Functional enrichment analysis was performed using Gene Ontology (GO) terms through g:Profiler to identify biological processes and signaling pathways associated with the proteins included in the network^30^. Significantly enriched terms were identified using the default statistical settings of the platform, and pathways relevant to diabetic neuropathy were selected for visualization.

### Validation of hub genes in a human dataset

A human DRG gene expression dataset obtained from dbGaP (accession phs002548.v1.p1)^31^ was used to validate the expression patterns of the selected hub genes. Receiver operating characteristic (ROC) curve analysis was performed to evaluate the potential of the selected hub genes as biomarkers of early neuronal dysfunction associated with diabetic neuropathy. The area under the curve (AUC) was calculated to assess the ability of each gene to discriminate between diabetic and control samples. Statistical significance was determined by testing whether the AUC differed significantly from 0.5, with p < 0.05 considered statistically significant.

### Statistical analysis

Statistical analyses were conducted in GraphPad Prism 9. For the frequency of F-waves occurrence within a 20-stimulus train, any potential exceeding 0.30 mV within the expected latency window was considered an F-wave response. Survival data were analyzed via the Log-rank test and visualized using the Kaplan-Meier plot. The significance threshold was set at p < 0.05. Additional statistical tests are indicated in the figure caption where appropriate.

## Ethics statement

All animal procedures were conducted in accordance with the institutional guidelines of the Center for Genetic Engineering and Biotechnology (CIGB) and complied with national and international regulations for the care and use of laboratory animals, including the Guide for the Care and Use of Laboratory Animals (8th edition, National Research Council, 2011).

## Consent for publication

Not applicable

## Availability of data and materials

Additional data are available from the corresponding author upon reasonable request.

## Authors’ contributions

G.R.H. conceived the study, performed the experiments and bioinformatic analyses, analyzed the data, prepared the figures, and wrote the manuscript. M.E.V.T. contributed to manuscript review and editing. All authors read and approved the final manuscript.

## Acknowledgments

The author thanks Dr. Héctor Pérez-Saad, MSc.Daniela Risco-Acevedo, Dr. Diana García del Barco, and Dr. Maday Fernández Mayola for their valuable support throughout the project.

